# Modeling structure, stability and flexibility of double-stranded RNAs in salt solutions

**DOI:** 10.1101/332676

**Authors:** L. Jin, Y.Z. Shi, C.J. Feng, Y.L. Tan, Z.J. Tan

**Author notes:** The authors contributed equally to the work (L. Jin, Y.Z. Shi).

## Abstract

Double-stranded (ds) RNAs play essential roles in many processes of cell metabolism. The knowledge of three-dimensional (3D) structure, stability and flexibility of dsRNAs in salt solutions is important for understanding their biological functions. In this work, we further developed our previously proposed coarse-grained model to predict 3D structure, stability and flexibility for dsRNAs in monovalent and divalent ion solutions through involving an implicit structure-based electrostatic potential. The model can make reliable predictions for 3D structures of extensive dsRNAs with/without bulge/internal loops from their sequences, and the involvement of the structure-based electrostatic potential and corresponding ion condition can improve the predictions on 3D structures of dsRNAs in ion solutions. Furthermore, the model can make good predictions on thermal stability for extensive dsRNAs over the wide range of monovalent/divalent ion concentrations, and our analyses show that thermally unfolding pathway of a dsRNA is generally dependent on its length as well as its sequence. In addition, the model was employed to examine the salt-dependent flexibility of a dsRNA helix and the calculated salt-dependent persistence lengths are in good accordance with experiments.

## Introduction

RNAs play a pervasive role in gene regulation and expression. In addition to single-stranded (ss) RNAs such as mRNAs and tRNAs, double-stranded (ds) RNAs are widespread in cells and are involved in a variety of biological functions (1–3). For examples, small noncoding dsRNAs can play a critical role in mediating neuronal differentiation (4); dsRNA segments of special lengths can inhibit the translation of mRNA molecules into proteins through attaching to mRNAs (5,6); and dsRNAs of more than 30 base-pair (bp) length can be key activators of the innate immune response against viral infections (7). Generally, dsRNAs realize their biological functions through becoming partially melted or changing their conformations (2-9). Furthermore, the inter-chain interactions in stabilizing structures of dsRNAs are very sensitive to the environment (e.g., temperature and ion conditions) (10-14). Thus, a full understanding of dsRNA-mediated biology would require the knowledge of three-dimensional (3D) structures, structural stability and flexibility of dsRNAs in ion solutions.

The 3D structures of RNAs including dsRNAs can be measured by several experimental methods such as X-ray crystallography, nuclear magnetic resonance (NMR) spectroscopy, and cryo-electron microscopy. However, it is still technically challenging and expensive to experimentally derive 3D structures of RNAs at high resolution and the RNA structures deposited in Protein Data Bank (PDB) are still limited (15). Therefore, as complementary methods, some computational models have been developed in recent years, aiming to predict RNA 3D structures in silico (16-23). The fragment assembly models (24-31) such as MC-Fold/MC-Sym pipeline (24), 3dRNA (25-27), RNAComposer (28) and Vfold3D (29,30) can successfully predict 3D structures of RNAs including even large RNAs at fast speed, however, these methods are generally based on given secondary structures and the limited known RNA 3D structures deposited in PDB. Although the fragment assembly method of FARNA (31) can predict 3D structures for RNAs from sequences, it could be only efficient for small RNAs due to its full-atomic resolution. In parallel ways, some coarse-grained (CG) models (32-41) such as iFold (42), SimRNA (43), HiRE-RNA (44) and RACER (45,46) have been proposed to predict 3D structures for RNAs with medium-lengths from their sequences based on knowledge-based statistical potentials or/and experiential parameters. However, these existing 3D structure prediction models seldom make quantitative predictions for thermodynamic stability and flexibility of RNAs.

Simultaneously, some models have been employed to predict thermodynamics of RNAs. Vfold2D/Vfold Thermal (29,30) with involving thermodynamic parameters can make reliable predictions on the free energy landscape of RNAs including pseudoknots at secondary structure level. The model proposed by Denesyuk and Thirumalai (47,48) can well predict the thermodynamics of small RNAs, while such structure-based (Gö-like) model could not predict 3D structures of RNAs solely from the sequences. Although other models such as iFold (42), HiRE-RNA (44), oxRNA (49) and NARES-2P (50,51) may give melting curves of RNAs, there is still lack of extensive experimental validation for these models.

Furthermore, RNAs are highly charged polyanionic molecules, and RNA structure and stability are generally sensitive to solution ion conditions, especially multivalent ions such as Mg^2+^ (8,10-14). The role of ions in RNA structure and stability, especially the role of Mg^2+^ which is generally beyond the mean-field descriptions (52,53), is seldom involved in the existing 3D structure prediction models. To predict the 3D structures and stability of RNAs in ion solutions, we have developed a CG model with implicit electrostatic potential (54,55), and the model has been validated through making reliable predictions on 3D structures and stability of RNA hairpins and pseudoknots as well as the ion effect on their stability. However, the model at present version is only applicable for ssRNAs and the implicit electrostatic potential of ions is independent on RNA structures. Thus, the model cannot make reliable predictions on 3D structure and stability for dsRNAs and may not detailedly predict the ion effect on 3D structures of RNAs.

In this work, we further developed our previous three-bead CG model for ssRNAs to predict 3D structures and stability of dsRNAs from their sequences. Furthermore, in the new version of the model, an implicit structure-based electrostatic potential is introduced in order to capture the effect of ions such as Mg^2+^ on 3D structures and stability of dsRNAs. As compared with the extensive experimental data, the present model can predict the 3D structures, stability and flexibility of various dsRNAs with high accuracy, and the effects of monovalent/divalent ions on the stability and flexibility of dsRNAs can be well captured by the present model. Additionally, our further analyses show that thermally unfolding pathway of a dsRNA is dependent on not only its length but also its sequence.

## Model and methods

### Coarse-grained structure model and energy function

To reduce the complexity of nucleotides, in our CG model, one nucleotide is represented by three beads: phosphate bead (P), sugar ring bead (C) and base bead (N) (54,55). The P and C beads are placed at the P and C4’ atom positions, and the base bead (N) is placed at N9 atom position for purine or N1 for pyrimidine; see Fig. 1. The three beads are treated as van der Waals spheres with the radii of 1.9 Å, 1.7 Å and 2.2 Å, respectively (54,55).

**FIGURE 1.**
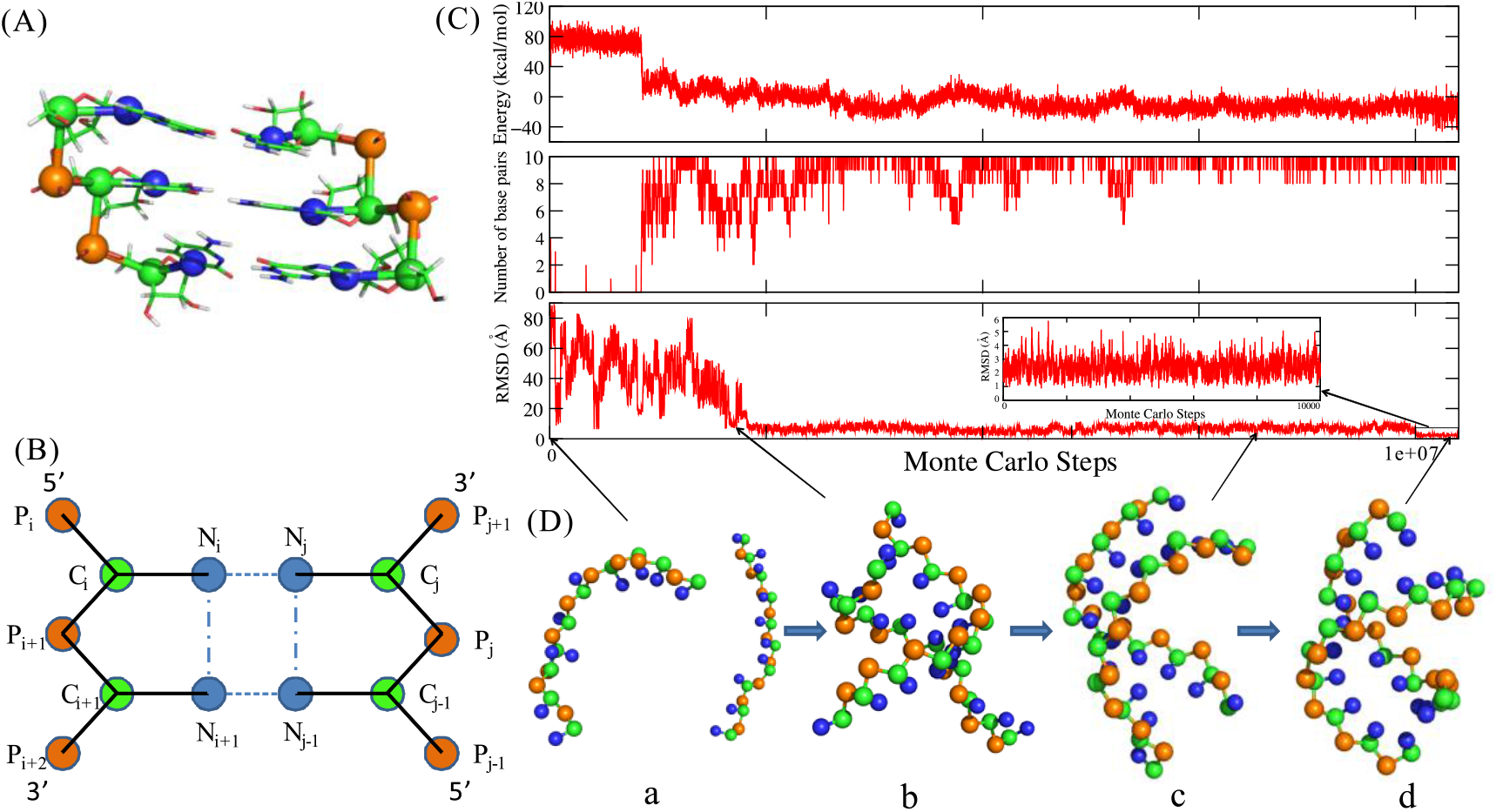
(A) The coarse-grained representation for one fragment of a dsRNA superposed on an all-atom representation. The beads of P (orange) and C (green) are located at P atom in phosphate group and C4’ atom in sugar ring, respectively. The beads of N (blue) are located at N9 atom position for purine or N1 atom position for pyrimidine. (B) The schematic representation for pseudo bonds (solid lines), base pairing (dashed lines) and base stacking (dash-dotted line) in the present model. (C, D) Illustration for the folding process of a typical dsRNA (PDB code: 2jxq) in the present model. (C) The system energy (top panel), number of base pairs (middle panel) and the RMSD (bottom panel) along the simulated annealing MC simulation. The inserted panel is the zoomed RMSD of the structure in the refinement procedure at the end of the simulation. (D) The four typical conformation states in the folding procedure. The structures are shown with PyMol (http://www.pymol.org).

The potential energy of a CG dsRNA is composed of two parts, bonded potential *U_bonded_* and *U_nonbonded_* potential (54,55)

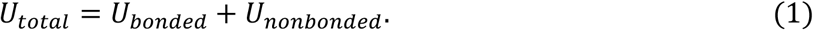

The bonded potential *U_bonded_* represents the energy associated with pseudo-covalent bonds between contiguous CG beads within any single chain, which includes bond length energy *U_b_*, bond angle energy and dihedral angle energy *U_d_*:

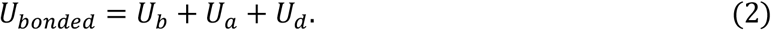

The initial parameters of these potentials were derived from the statistical analysis on the available 3D structures of RNA molecules in PDB (http://www.rcsb.org/pdb/home/home.do), and two sets of parameters *Para_helical_* and *Para_nonhelical_* were provided for stems and single strands/loops, respectively. Note that only *Para_nonhelical_* is used in folding process and both of *Para_helical_* and *Para_nonhelical_* are used in structure refinement (54,55). The nonbonded potential *U _nonbonded_* in Eq. 1 includes the following five components

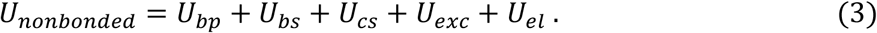

*U_bp_* is the base-pairing interaction between Watson-Crick (G-C and A-U) and wobble (G-U) base pairs. *U_bs_* and *U_cs_* are sequence-dependent base stacking and coaxial stacking interactions between two neighbour base pairs and between two neighbour stems, respectively. The strengths of *U_bs_* and *U_cs_* were derived from the combined analysis of available thermodynamic parameters and Monte Carlo (MC) simulations (54,55). *U_exc_* represents the excluded volume interaction between two CG beads and it is modelled by a purely repulsive Lennard-Jones potential.

The last term *U_el_* in Eq. 3 is a structure-based electrostatic energy for an RNA, which is newly refined in the model to better capture the contribution of monovalent and divalent ions to RNA 3D structures. The electrostatic potential is treated as a combination of Debye-Hückel approximation and the counterion condensation (CC) theory (52-55)

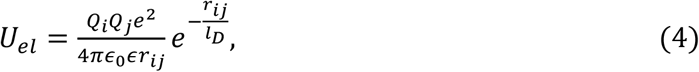

where *r_ij_* is the distance between the *i*-th and *j*-th P beads, each of which carries a unit negative charge (-*e*). *l_D_* is the Debye length of ion solution. *ϵ*_0_ is the permittivity vacuum and *ε* is the effective temperature-dependent dielectric constant of water (54,55). The reduced negative charge *Q_i_* on the *i*-th P bead is given by

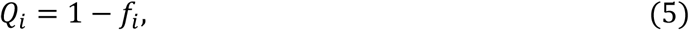

where *f_i_* is the fraction of ion neutralization. In the present model, beyond the assumption of uniform distribution of binding ions along RNA chain in our previous model, *f_i_* is dependent on RNA 3D structure, and includes the contributions of monovalent and divalent ions

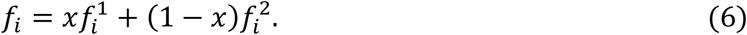

Here, *x* and 1 – *x* represent the contribution fractions from monovalent and divalent ions, which can be derived from the tightly bound ion (TBI) model (55-60). 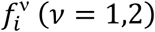 is the binding fraction of *v*-valent ions, and is given by

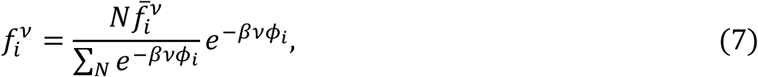

where *N* is the number of P beads. 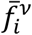 represents the average charge neutralization fraction of ions and the CC theory gives that (52-55) 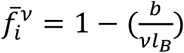, where *b* is the average charge spacing on RNA backbone and *l_B_* is Bjerrum length. *ϕ_i_* in Eq. 7 is the electrostatic potential at *i*-th P bead and can be approximately calculated by

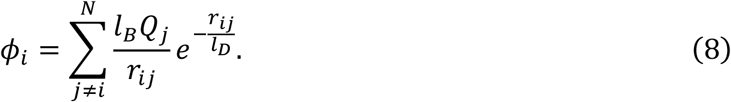

Eqs. 5-8 show that the structure-based reduced charge fraction *Q_i_* needs to be obtained through an iteration process; see more details in Supporting Material.

The detailed descriptions on the CG energy function as well as the parameters for the potentials in Eqs. 1-3 can be found in the Supporting Material.

### Simulation algorithm

To effectively avoid the traps in local energy minima, the MC simulated annealing algorithm is used to sample conformations for a dsRNA at given monovalent/divalent ion conditions. Based on the sequence of a dsRNA, two initial random CG chains can be generated and be separately placed in a cubic box, the size of which is determined by concentration of ssRNA. Generally, the simulation of a dsRNA system with a given ion condition is performed from a high temperature (e.g., 110°C) to the target temperature (e.g., room/body temperature). At each temperature, the conformations of the dsRNA are sampled by intra-strand pivot moves and inter-strand translation/rotation through the Metropolis algorithm until the system reaches enough equilibrium. In this process, the newly refined electrostatic potential *U_el_* is involved (see Eq. 4), and *U_el_* can only be obtained after an iterative process for *Q_i_*. In practice, *U_el_* is renewed over every 20 MC steps, and generally, *U_el_* can be obtained through ~4 times iterations for converged *Q_i_*. Thus the increase in computation cost due to involving the newly refined *U_el_* is negligible compared to the whole simulation cost. The equilibrium conformations of the system at each temperature can be saved to obtain 3D structures and structural properties of the dsRNA at each temperature.

### Calculation of melting temperature

The stability of dsRNAs generally depends on strand concentration due to the contribution of translation entropy of melted ssRNA chains (61). However, for a dsRNA with low strand concentrations (e.g., 0.1 mM in experiments), a very long simulation time is generally required to reach equilibrium for the dsRNA system. To make our calculations efficient, the simulations for dsRNAs can be performed at relatively high strand concentrations 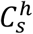 (e.g., ~10 mM for dsRNAs with length ≤10-bp and ~1 mM for dsRNAs with length >10-bp) (44,62). Based on the equilibrium conformations at each temperature *T*, the fraction Φ(*T*) of unfolded state characterized as completely dissociated single-stranded chain can be obtained at *T*. Since the small system of the simulation (two strands in a simulational box) can lead to a significant finite-size effect (63), the predicted Φ(*T*) needs to be further corrected to the fraction *f_h_*(*T*) of unfolded state at the high bulk strand concentration 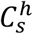 (63):

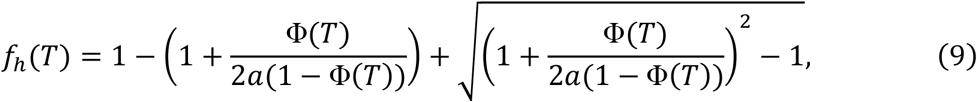

where *a*=1 and 2 for nonself-complementary and self-complementary sequences, respectively (63). Afterwards, based *f_h_*(*T*) on at the high strand concentration, the fraction of unfolded state *f*(*T*) at an experimental strand concentration *C_s_* (e.g., ~0.1 mM) can be calculated by (64,65)

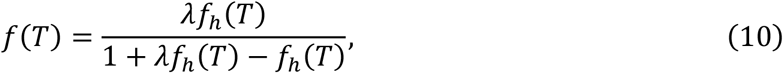

where 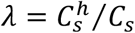. Finally, the fractions of unfolded state *f*(*T*) can be fitted to a two-state model to obtain the melting temperature *T_m_*(54,55,64),

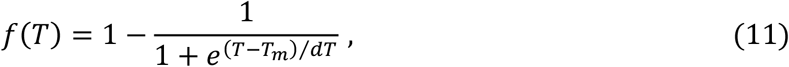

where *dT* is an adjustable parameter. More details about the calculation of melting temperature are given in the Supporting Material. For long dsRNAs whose unfolding can be non-two-state transition, we still used the above formulas to estimate their melting temperatures, in analogy to related experiment (66).

## Results and discussion

In this section, the present model was first employed to predict 3D structures of extensive dsRNAs in monovalent/divalent ion solutions. Afterwards, the model was used to predict stability of extensive dsRNAs and the effects of monovalent/divalent ions, and further to analyze the thermally unfolding pathway of various dsRNAs. Finally, the model was employed to examine the salt-dependent flexibility of a dsRNA helix. Our predictions and analyses were extensively compared with the available experiments and existing models.

### Structure predictions for dsRNAs in ion solutions

Two sets of available dsRNAs were used in this work on 3D structure prediction. One set includes 16 dsRNAs whose structures were determined by X-ray experiments (defined as X-ray set), and the other set contains 10 dsRNAs whose structures were determined by NMR experiments in ion solutions (defined as NMR set). The PDB codes as well as the descriptions of the dsRNAs in two sets are shown in Tables 1 and S3, respectively. In the following, we first made predictions for the 26 structures of dsRNAs in X-ray set and NMR set at high salt concentration (e.g., 1 M Na^+^). Afterwards, we made predictions for the 3D structures of dsRNAs in NMR set at respective ion conditions.

#### Structure predictions for dsRNAs at 1M [Na^+^]

For 26 dsRNAs in X-ray set and NMR set, the 3D structures were predicted from sequences with strand concentration of 0.1 mM at high salt concentration (e.g., 1 M Na^+^), regardless of possible ion effects. In the following, we used a paradigm dsRNA (PDB code: 2jxq; shown in Table 1) to show the structure predicting process of dsRNA with the present model, which is shown in Fig. 1C. First, the energy of the system reduces with the decrease of temperature (from 100°C to room temperature) and the dsRNA folds into native-like structures (e.g., structure c in Fig. 1D) from an initial random configuration (e.g., structure a in Fig. 1D). Second, a further structure refinement (~ 1.2 × 1 0 ^7^ MC steps) is performed at the target temperature (e.g., room temperature), where the last predicted structure from the annealing process is taken as input and the parameters *Para*_nonhelical_ of bonded potentials are replaced by *Para*_helical_ for the base-paired regions in order to better capture the geometry of helical stems (54,55). Finally, an ensemble of refined 3D structures (~10000 structures) can be obtained over the last ~ MC steps, and these structures can be evaluated by the root-mean-square deviation (RMSD) values calculated over all the beads in predicted structures from the corresponding atoms in the native structures in PDB (67). As shown in Fig. 1C and Fig. 2, for the dsRNA of PDB code 2jxq, the mean RMSD (the averaged value over the refined structures) and the minimum RMSD (from the structure closest to the native one) are 2.1 Å and 0.8 Å, respectively.

**FIGURE 2.**
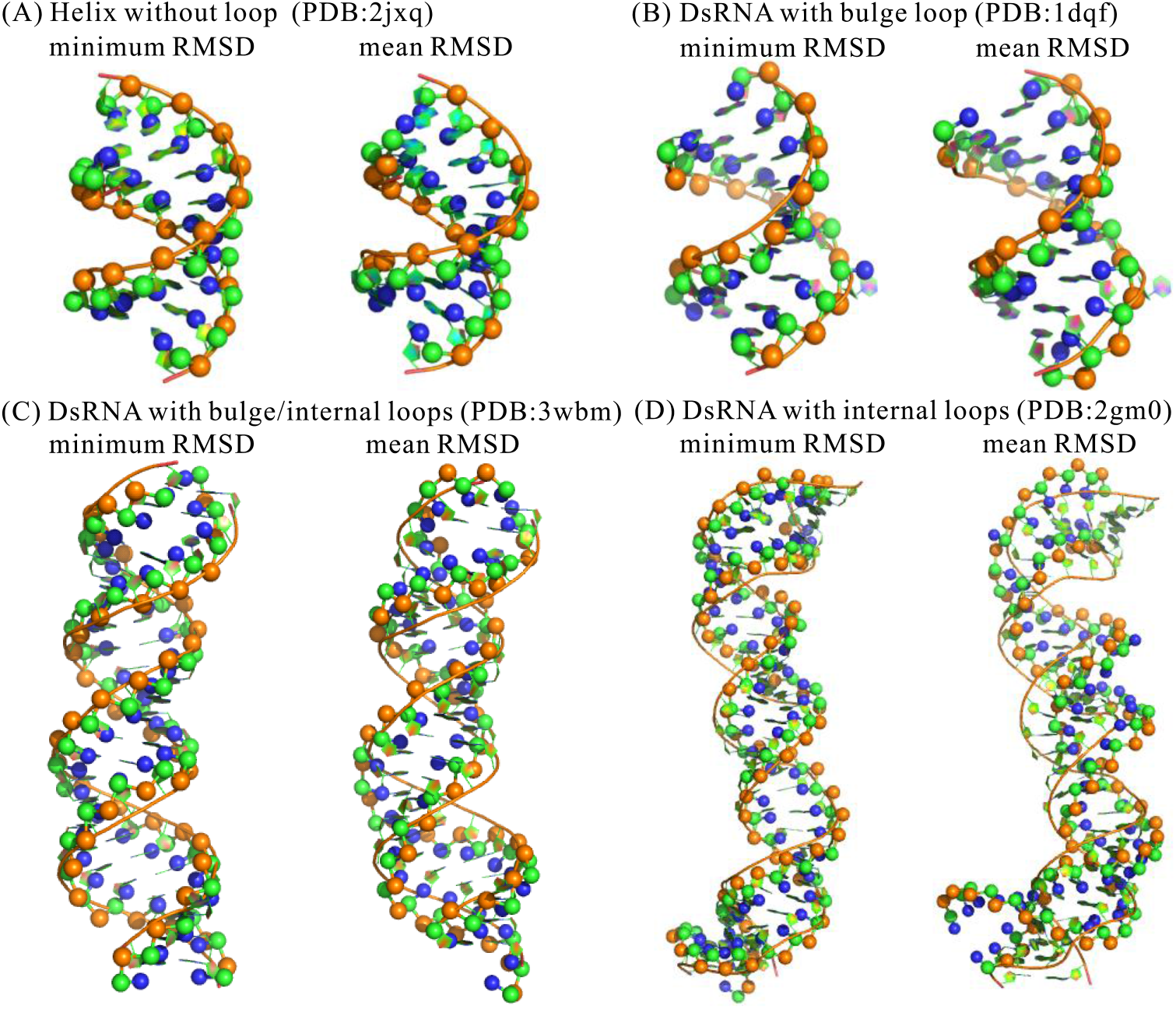
The predicted 3D structures (ball-stick) with mean/minimum RMSDs, in comparison with the corresponding native structures (cartoon) for four typical dsRNAs. (A) A dsRNA helix without loop (PDB code: 2jxq) with mean/minimum RMSDs of 2.0 Å/0.8 Å. (B) A dsRNA containing a bulge loop (PDB code: 1dqf) with mean/minimum RMSDs of 2.5 Å/1.6 Å. (C) A dsRNA containing 2 bulge loops and 2 internal loops (PDB code: 3wbm) with mean/minimum RMSDs of 5.4 Å/2.3 Å. (D) A dsRNA containing 4 internal loops (PDB code: 2gm0) with mean/minimum RMSDs of 6.1 Å/3.1 Å. The structures are shown with PyMol (http://www.pymol.org).

**Table 1.**
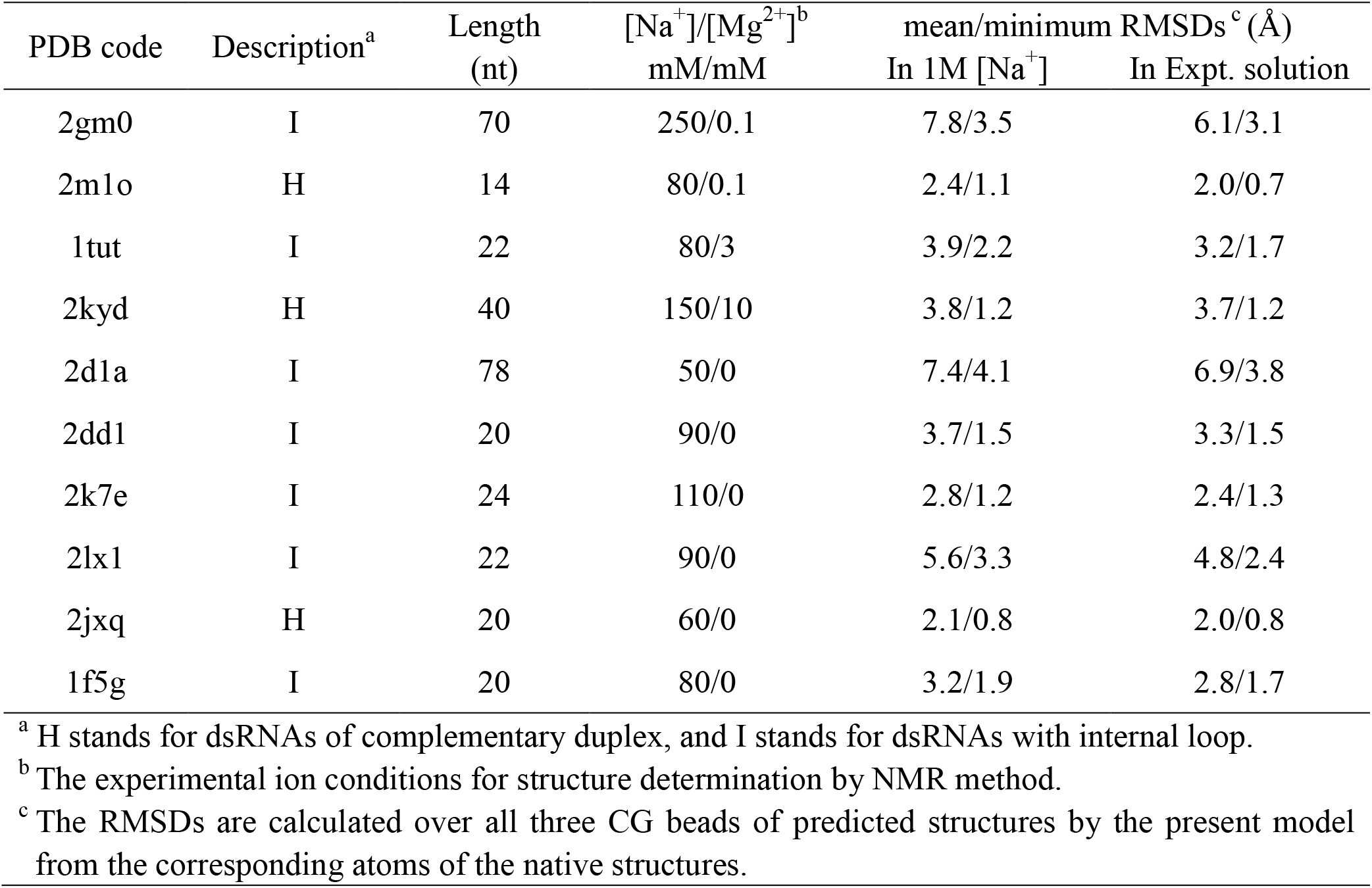
10 dsRNAs in NMR set for structure prediction at respective salt conditions.

| PDB code | Description<sup>a</sup> | Length<br>(nt) | [Na<sup>+</sup>]/[Mg<sup>2+</sup>]<sup>b</sup><br>mM/mM | mean/minimum RMSDs<sup>c</sup> (Å) |  |
| --- | --- | --- | --- | --- | --- |
|  |  |  |  | In 1M [Na<sup>+</sup>] | In Expt. solution |
| 2gm0 | I | 70 | 250/0.1 | 7.8/3.5 | 6.1/3.1 |
| 2m1o | H | 14 | 80/0.1 | 2.4/1.1 | 2.0/0.7 |
| 1tut | I | 22 | 80/3 | 3.9/2.2 | 3.2/1.7 |
| 2kyd | H | 40 | 150/10 | 3.8/1.2 | 3.7/1.2 |
| 2d1a | I | 78 | 50/0 | 7.4/4.1 | 6.9/3.8 |
| 2dd1 | I | 20 | 90/0 | 3.7/1.5 | 3.3/1.5 |
| 2k7e | I | 24 | 110/0 | 2.8/1.2 | 2.4/1.3 |
| 2lx1 | I | 22 | 90/0 | 5.6/3.3 | 4.8/2.4 |
| 2jxq | H | 20 | 60/0 | 2.1/0.8 | 2.0/0.8 |
| 1f5g | I | 20 | 80/0 | 3.2/1.9 | 2.8/1.7 |

<sup>a</sup> H stands for dsRNAs of complementary duplex, and I stands for dsRNAs with internal loop.
<sup>b</sup> The experimental ion conditions for structure determination by NMR method.
<sup>c</sup> The RMSDs are calculated over all three CG beads of predicted structures by the present model from the corresponding atoms of the native structures.

Following the above process, the 3D structures of 26 dsRNAs including 15 dsRNAs with bulge/internal loops were predicted by the present model with the overall mean RMSD of ~3.3 Å and the overall minimum RMSD of ~1.8 Å; see Figs. 2 and 3 and Table S1 in the Supporting Material. This shows that the present model with coaxial/base stacking can reliably capture the 3D shapes of various dsRNAs including those with bulge/internal loops. For 13 dsRNAs with internal loops, the overall mean RMSD is ~4.2 Å, which is slightly larger than that (~3.3 Å) of all the 26 dsRNAs. This is because large internal loops generally contain non-canonical base pairs, which is ignored in the present model. For example, the dsRNA of PDB code 3wbm contains 2 internal loops with several non-canonical base pairs to keep the helix more continuous than the predicted one; see Fig. 2C.

**FIGURE 3.**
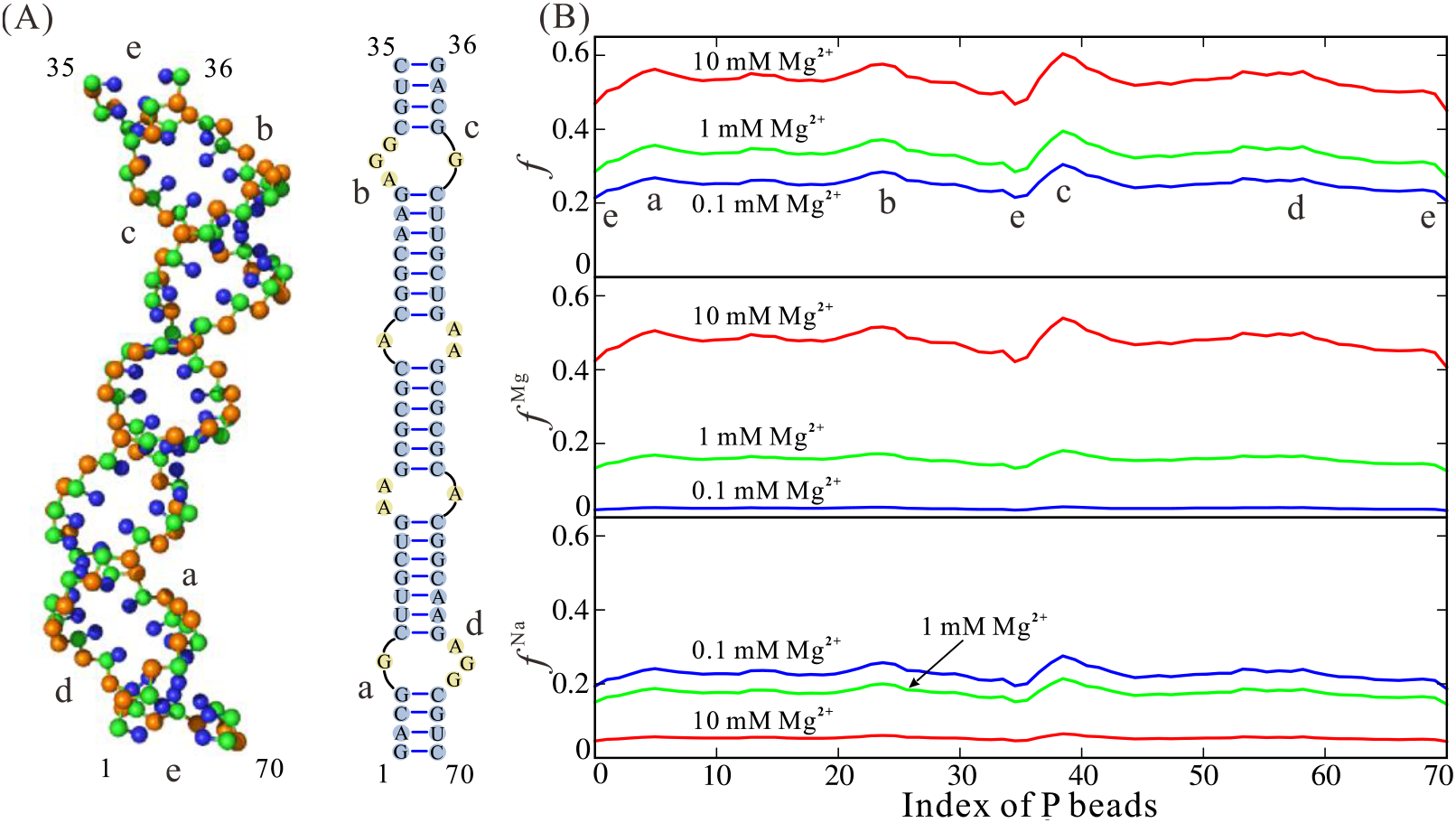
(A) The predicted 3D structure (left, shown with PyMol) and secondary structure (right) of a dsRNA (PDB: 2gm0). (B) The calculated ion charge neutralization fractions along P beads of the predicted structure of the dsRNA at different ion conditions; see Eq. 6. *f*: the total ion neutralization fractions (top panel). *f*_Mg_: neutralization fractions of Mg^2+^ (middle panel). *f*_Na_: neutralization fractions of Na^+^ (bottom panel). The blue, green and red lines denote the cases of the dsRNA in 150 mM Na solutions mixed with 0.1 mM, 1 mM and 10 mM Mg^2+^, respectively. The two bent regions labeled with (a, d) and (b, c) induced by the internal loops correspond to the ion neutralization fraction peaks in panel B, and two helical ends of the dsRNA labeled with e correspond to the ion neutralization fraction troughs in panel B.

#### Structure predictions for dsRNAs in respective ion solutions

Since RNA structures can be strongly influenced by ions (10-14), we introduced the structure-based electrostatic potential in the present model to improve the 3D structure prediction for dsRNAs at the respective ion conditions. In the following, we first examined the structure-based electrostatic potential through the charge neutralization fractions *f*^Na^ and *f*^Mg^ of Na^+^ and Mg^2+^ along an example dsRNA (PDB code: 2gm0) in mixed Na^+^/Mg^2+^ solutions. As shown in Fig. 3B, *f*^Na^ and *f*^Mg^ appear dependent on Na^+^/Mg^2^+ concentrations as well as dsRNA structures: (i) as Mg^2+^ concentration increases, increases and decreases, due to the competition between Mg^2+^ and Na^+^ in binding to an RNA and lower binding entropy penalty for Mg^2+^ at higher Mg^2+^ concentration (10-14,56-60); (ii) *f*^Na^ and *f*^Mg^ are larger at bent regions and appear less at two ends, which is attributed to the higher P beads charge density at bending regions and lower P beads charge density at two ends of the dsRNA. Therefore, the newly refined electrostatic potential (Eqs. 4-8) can capture the structure-based ion binding and the competitive binding between Na^+^ ions and Mg^2+^ ions to dsRNAs.

To examine whether the involvement of the implicit structure-based electrostatic potential (Eqs. 4-8) and corresponding ion conditions can improve 3D structure prediction for dsRNAs, we further predicted the 3D structures for 10 dsRNAs in NMR set at their respective experimental ion conditions. For the 10 dsRNAs, the overall mean RMSD between predicted structures in ion solutions and the native structures is ~3.7 Å, which is visibly smaller than that (~4.3 Å) of the predictions at 1 M [Na^+^]; see Table 1. This suggests that the inclusion of structure-based electrostatic potential and corresponding ion conditions in this model can improve the predictions on 3D structures for dsRNAs in ion solutions. Furthermore, Table 1 also shows that such improvement appears more pronounced for the dsRNAs with internal loops and for the ion conditions containing Mg^2^+, e.g., mean RMSDs of the dsRNA of 2gm0 and 1tut decrease from 7.8 Å and 3.9 Å to 6.1 Å and 3.2 Å, respectively. This indicates that the newly refined electrostatic potential can effectively involve the RNA structure information and the effect of ions such as Mg^2+^.

#### Comparisons with other models

To further examine the present model, we made extensive comparisons with three existing RNA structure prediction models: FARNA (31), RACER (45,46) and MC-Fold/MC-Sym pipeline (24). FARNA is a fragment-assembly model with high resolution for small RNAs (31). As shown in Fig. 4A, the average mean RMSD (~3.9 Å) from the present model is very slightly smaller than that (~4.1 Å) from FARNA. Afterwards, we made the comparison with a newly developed CG model RACER (45). As shown in Fig. 4B, the mean RMSD of the predictions from our model (~2.6 Å) is slightly smaller than that (~3.2 Å) from RACER. Furthermore, we made the extensive comparison with MC-Fold/MC-Sym pipeline (24), a well-established RNA 2D/3D structure prediction model with a web server (http://www.major.iric.ca/MC-Pipeline). We used the web server of MC-Fold/MC-Sym pipeline to predict the structures of all the dsRNAs involved in our structure prediction and chose the top 1 predicted structure to make comparisons with the present model. As shown in Fig. 4C, the average mean RMSD (~3.3 Å) of the predictions from the present model is slightly smaller than that (~3.8 Å) of the top 1 structures from MC-Fold/MC-Sym pipeline. Therefore, the above comparisons show that the present model can be reliable in predicting 3D structures of dsRNAs. Beyond 3D structure predictions, the present model can also predict stability and flexibility of dsRNAs in ion solutions.

**FIGURE 4.**
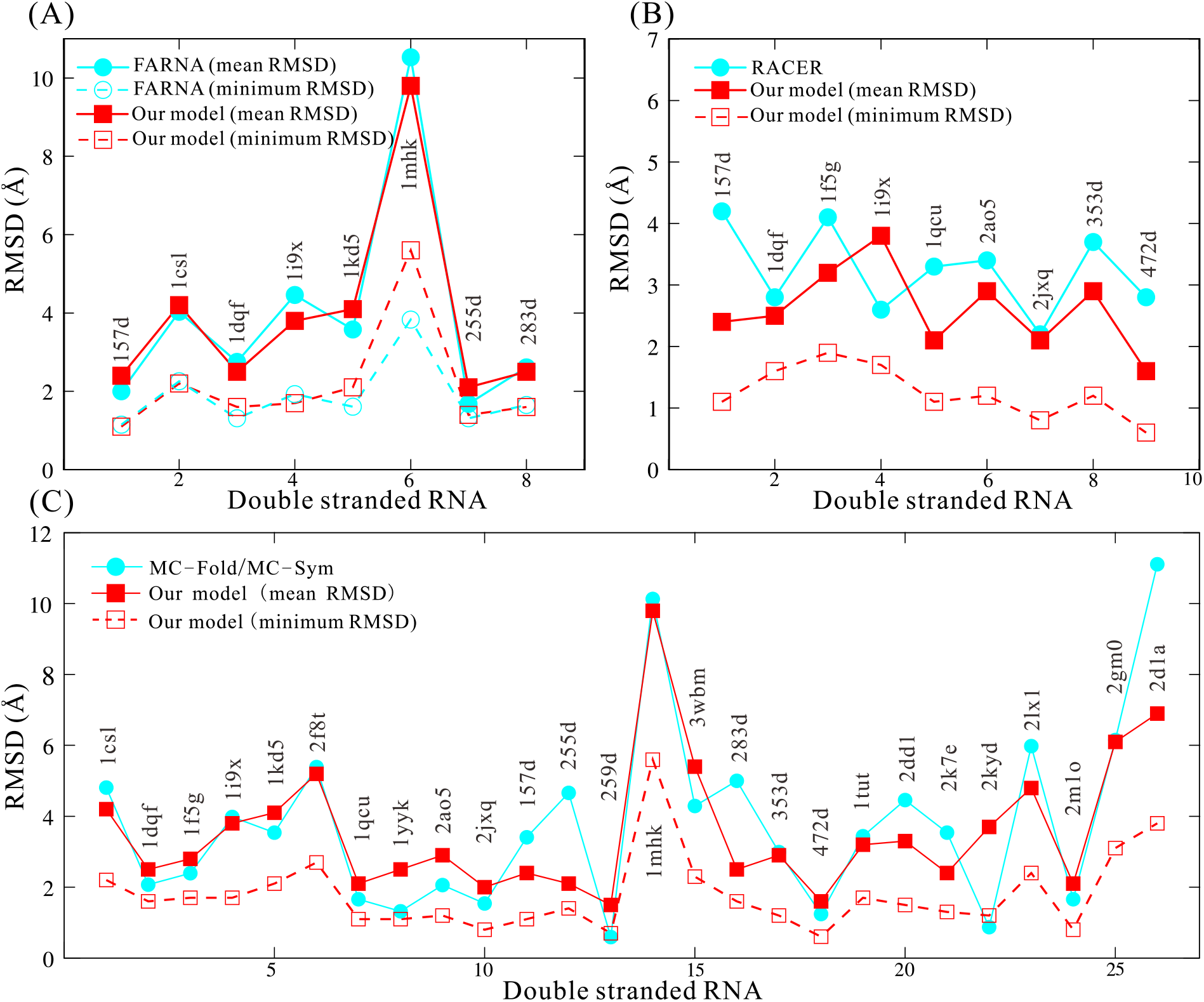
The comparisons of the predicted 3D structures between the present model and the existing models: (A) FARNA (31), (B) RACER (45,46) and (C) MC-Fold/MC-Sym pipeline (24). The RMSDs of structures predicted by FARNA are calculated over the C4’ atom (31). The RMSDs of structures predicted by RACER, MC-Fold/MC-Sym pipeline and the present model are calculated over all CG beads (24,45). The data of FARNA and RACER are taken from Ref. 31 and Ref. 45, respectively.

### Stability of dsRNAs in ion solutions

#### Stability of dsRNAs with various sequences

As described in the section of Model and methods, for a dsRNA with a given strand concentration, the melting curve as well as the melting temperature *T_m_* can be calculated by the present model. For example, for the sequence (CGCG)_2_, the melting curve of the dsRNA with a high strand concentration of 10 mM can be predicted based on the fractions of unfolded state at different temperatures, and the melting curve as well as the melting temperature *Tm* of the dsRNA at low experimental strand concentration (0.1 mM) can be calculated through Eqs. 9-11; see Figs. 5A and B. As shown in Figs. 5A and B, the predicted *T_m_* of the sample sequence (CGCG)_2_ with experimental strand concentration of 0.1 mM is ~19.5°C, which agrees well with the corresponding experimental value (~19.3° C). Furthermore, we further predicted the thermodynamic stability of 22 dsRNAs (4~14-bp) with various complementary sequences; see Table 2. Here, dsRNAs are assumed in solutions of 1 M [Na^+^], to solely examine the stabilities of dsRNAs of various sequence and make comparisons with extensive experimental data (65,66,68-70). As shown in Table 2, T_m_’s of extensive dsRNAs from the present model are in good agreement with the corresponding experimental data with the mean deviation ~1.3°C and maximum deviations <2.5°C. Such agreement indicates that the sequence-dependent base pairing and base stacking interactions in the present model can well capture the stability of dsRNAs of extensive sequences and different lengths (65,66,68-70).

**FIGURE 5.**
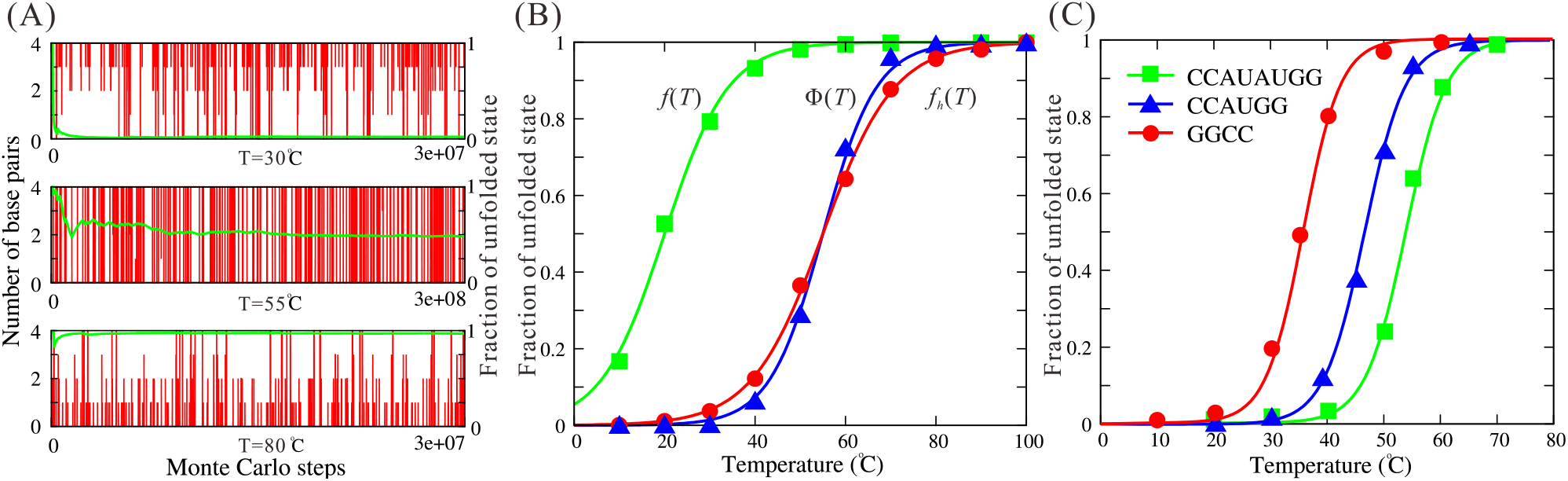
(A) The time-evolution of the number of base pairs (vertical or red lines) and the average fractions of unfolded state (transverse or green lines) for the sample dsRNA of (CGCG)2 at 30C (top panel), 55°C (middle panel) and 80° C (bottom panel). (B) The fractions of unfolded state as functions of temperature for the dsRNA of (CGCG)_2_. Symbols: the predicted data at different temperatures; blue triangle: at high strand concentration (10 mM); red circle: the corrected value of *f_h_*(*T*) by Eq. 9; green square: at experimental strand concentration (0.1 mM) derived by Eq. 10. Lines: the fitted melting curve to the predicted data through Eq. 11. More details can be found in Supporting Material. (C) The fractions of unfolded states for three dsRNAs with 0.1 mM strand concentration as functions of temperature. Symbols are the predicted data and lines are the fitted curves for the three dsRNAs through Eq.11.

**Table 2.**
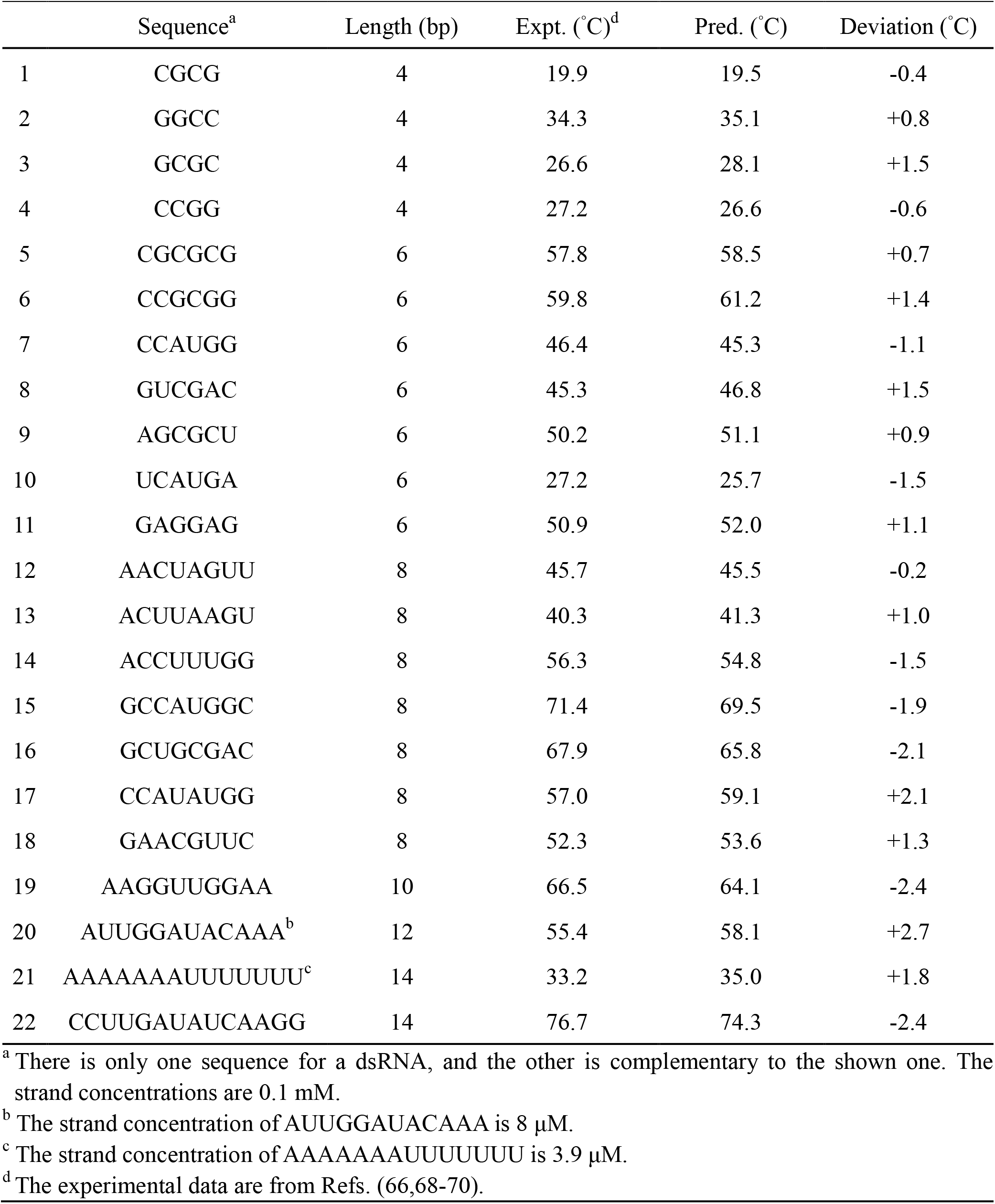
The melting temperatures *T_m_*’s for 20 dsRNAs in 1 M [Na_+_] solution.

#### Thermally unfolding pathways of dsRNAs

Since intermediate states of RNAs can be important to their functions (1-3,10,71,72), we made further analyses on thermally unfolding pathways for different dsRNAs. To distinguish the possible different states of dsRNAs at different temperatures in our simulations, all the states for a dsRNA were more detailedly divided into unfolded sate (U, two disassociated single strands), possible hairpin state (H, with at least one hairpin), folded helix state (F, with the formation of all base pairs except for the two end ones) and partially folded helix state (P, other conformations besides U, F and H states).

As shown in Fig. 6, the unfolding pathways of dsRNAs are dependent on their length as well as sequences. For short sequences (≤6-bp), dsRNAs undergo the standard two-state melting transitions and there is almost no intermediate states such as P and H states; see Figs. 6A and B, which is consistent with the previous experiments (66). As chain length increases to ~8-bp, P state begins to appear and can become visible at ~*T_m_*; see Figs. 6C and D. Figures 6C and D also show that the unfolding pathways of dsRNAs with the same chain length but different sequences would be slightly different, e.g., the fraction of P state of (AACUAGUU)_2_ with end A-U base pairs is slightly higher than that of (CCAUAUGG)_2_ with end G-C base pairs. This is because the unstable end A-U base pairs can induce more notable P state than stable end G-C base pairs; see Fig. S1 in the Supporting Material.

**FIGURE 6.**
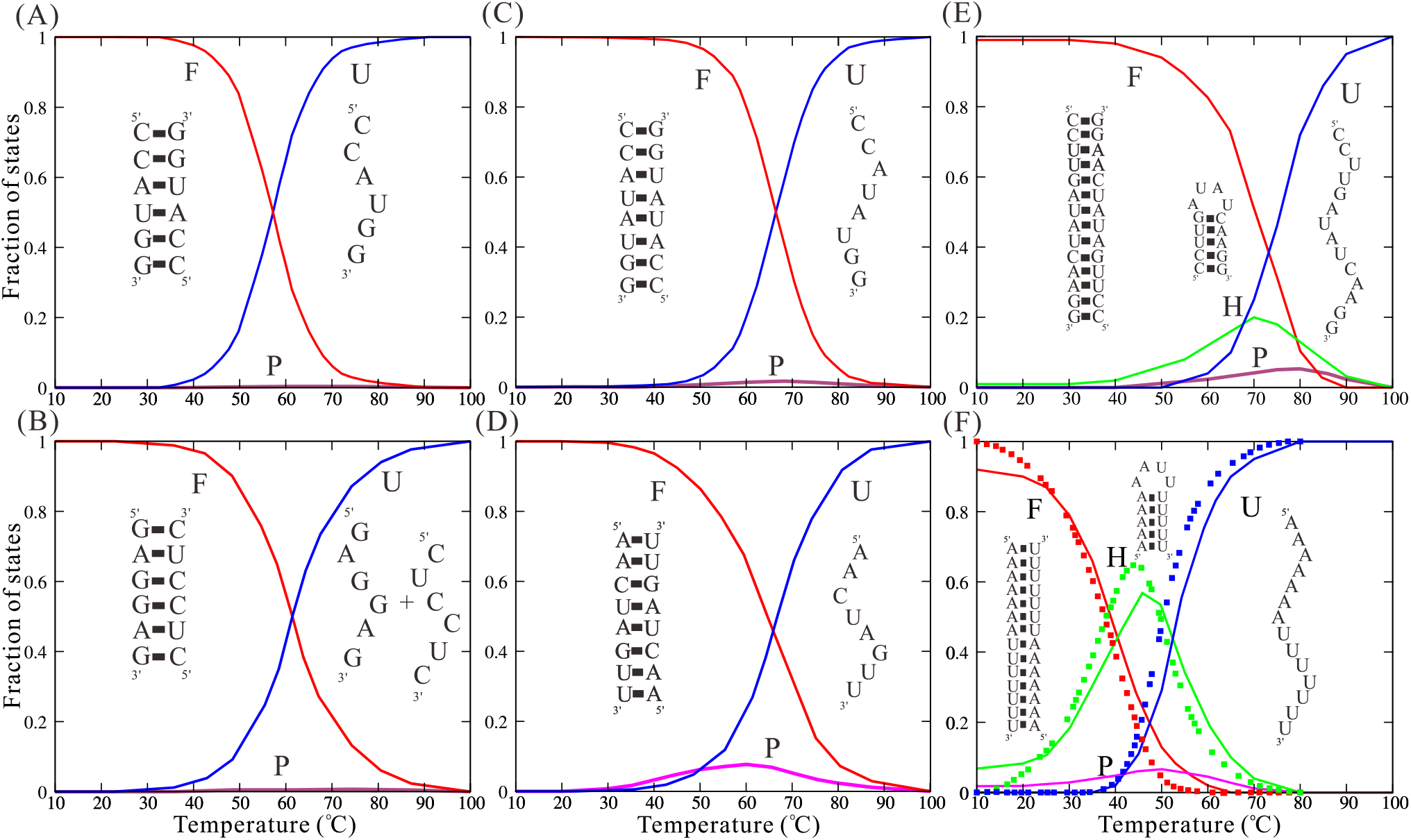
The fractions of F (Folded, red lines), P (Partially folded, pink lines), H (Hairpin, green lines) and U (Unfolded, blue lines) states as functions of temperature for the unfolding of (CCAUGG)_2_ (A), (GAGGAG,CUCCUC) (B), (CCAUAUGG)_2_ (C), (AACUAGUU)_2_ (D), (CCUUGAUAUCAAGG)_2_ (E) and (AAAAAAAUUUUUUU)_2_ (F). The full symbols in panel (F) are the correspondingly experimental data from Ref. 69. The insets in panels illustrate the typical predicted secondary structures for F, U and H states.

For dsRNAs with more than 10 base pairs, their thermally unfolding pathways become more complex and interesting, since their single-stranded chain may fold to hairpin structures. As shown in Figs. 6E and F for the dsRNA of (CCUUGAUAUCAAGG)_2_ and (AAAAAAAUUUUUUU)_2_, the fractions of F state are near unity at low temperature. As temperature increases, the dsRNAs begin to melt and H states would form from the melted ssRNAs with the maximum fractions of ~0.2 and ~0.5 for the two dsRNAs, respectively. At higher temperature, the dsRNAs almost become completely melted as U state. Notably, as shown in Fig. 6F, the unfolding pathway of (AAAAAAAUUUUUUU)_2_ predicted by the present model is very close to the corresponding experiments (66,69), suggesting that the melting pathways of dsRNAs can be well captured by our analyses with the present model. The difference on unfolding pathways between the two dsRNAs is attributed to the different sequences, i.e., G-C content, especially at two ends. Specifically, the fractions of states follow the order of F > *U* ≳ *H* > *P* for (CCUUGAUAUCAAGG)_2_ at ~70°C, while for (AAAAAAAUUUUUUU)_2_ at ~45° C, such order becomes H > *F* ≳ **U** > *P*. To reveal what determines the order, we calculated the stability for the states with Mfold (73). We found that the order of state fractions is in agreement with that of state stability. For example, the formation free energies for F, H and P states are ~-0.5 kcal/mol, ~0.2 kcal/mol and ~1.5 kcal/mol for (CCUUGAUAUCAAGG)_2_ at ~70°C, respectively. For (AAAAAAAUUUUUUU)_2_ at ~45° C, the formation free energies for H, F and P states are ~—0.8 kcal/mol, ~—0.1 kcal/mol and ~0.6 kcal/mol. This indicates that the unfolding pathway of dsRNA is dependent on the stability of possible states.

Although unfolding of long dsRNAs can be non-two-state transitions, in analogy to experiments (66), we can still estimate their melting temperatures by assuming strands completely disassociated state as U state (66); see Eq. 11 in section of Model and method.

#### Stability of dsRNAs with bulge/internal loops

Beyond the dsRNAs with complementary sequences shown above, the stability of other 8 dsRNAs with bulge/internal loops was examined by the present model. As shown in Table 3, for the dsRNAs with single/double bulge loops of different loop lengths (sequences 1-6) and the dsRNAs with internal loops (sequences 7 and 8), the mean deviation between the predicted *T_m_*’s and the experimental data (74-80) is ~2.6°C, which indicates that the present model with the coaxial stacking potential can roughly estimate the stability of dsRNAs with bulge/internal loops. However, such predictions especially for dsRNAs with long bulge/internal loops are not as precise as those for dsRNAs without loops. The detailed comparisons with experimental data show that, the predicted *T_m_*’s for the dsRNAs with 1 nucleotide (nt) bulge loop are slightly higher than experimental data, while the present model underestimates the stability of dsRNAs with longer bulge loops, which may suggest that the coaxial stacking potential *U_cs_* involved in the present model may slightly overestimate the coaxial interaction strength while underestimates the coaxial interaction range. For the dsRNAs with internal loop (e.g., AA/AA or AAA/AAA), the present model underestimates their stability, which may be attributed to the ignorance of non-canonical base pairs in the present model (80).

**Table 3.**
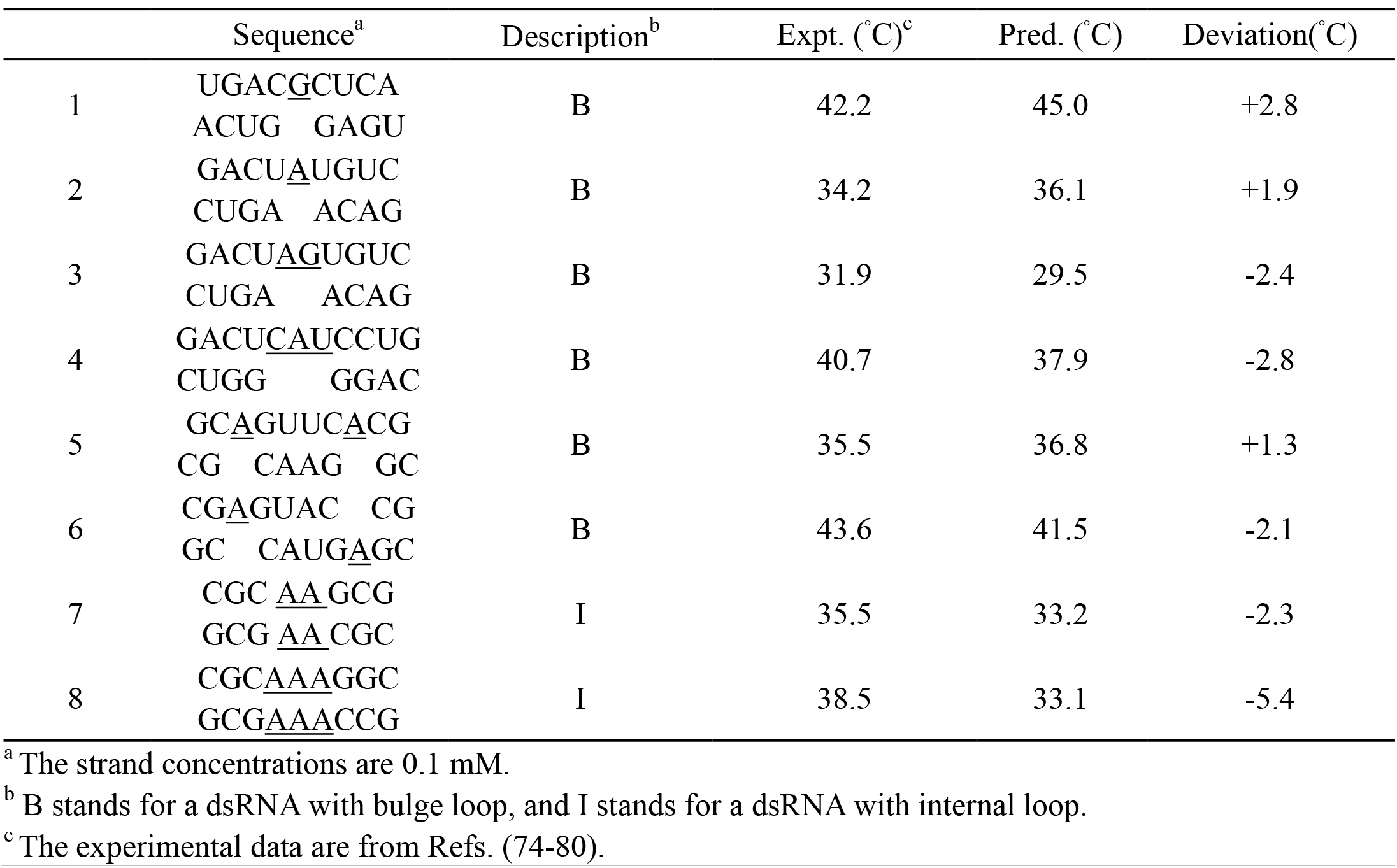
The melting temperatures *T_m_* for 8 dsRNAs with bulge/internal loop in 1 M Na^+^ solution.

#### Effects of monovalent and divalent ions

The thermal stability of RNA molecules is generally sensitive to the ionic conditions (10-14, 56-60). Particularly, Mg^2+^ is efficient in neutralizing the negative charges on RNA molecule and generally plays important role in RNA folding (14,56-60,81-84). However, most of the existing structure prediction models cannot quantitatively predict the stability of dsRNAs in ion solutions, especially in the presence of Mg^2+^. Here, we employed the present model to examine the stability for dsRNAs over a wide range of monovalent and divalent ion concentrations.

First, we examined the effect of monovalent ions on the stability of dsRNAs. As shown in Fig. 7A, for 5 dsRNAs with different sequences and lengths, the predicted melting temperatures *T_m_*’s from the present model agree well with the experimental data (65,66,68-70,84) with a mean deviation <2 °C over the wide range of [Na^+^]. As [Na^+^] increases from 10 mM to 1 M, *T_m_*’s of the dsRNAs increase obviously, which is attributed to lower ion-binding entropy penalty and stronger ion neutralization for base pair formation at higher [Na^+^]; see also Fig. S1 in the Supporting Material. Furthermore, Fig. 7A shows that the [Na^+^]-dependence of *T_m_* is stronger for longer dsRNAs. This is because base pair formation of longer dsRNAs causes larger build-up of negative charges and consequently causes stronger [Na^+^]-dependent ion-binding.

**FIGURE 7.**
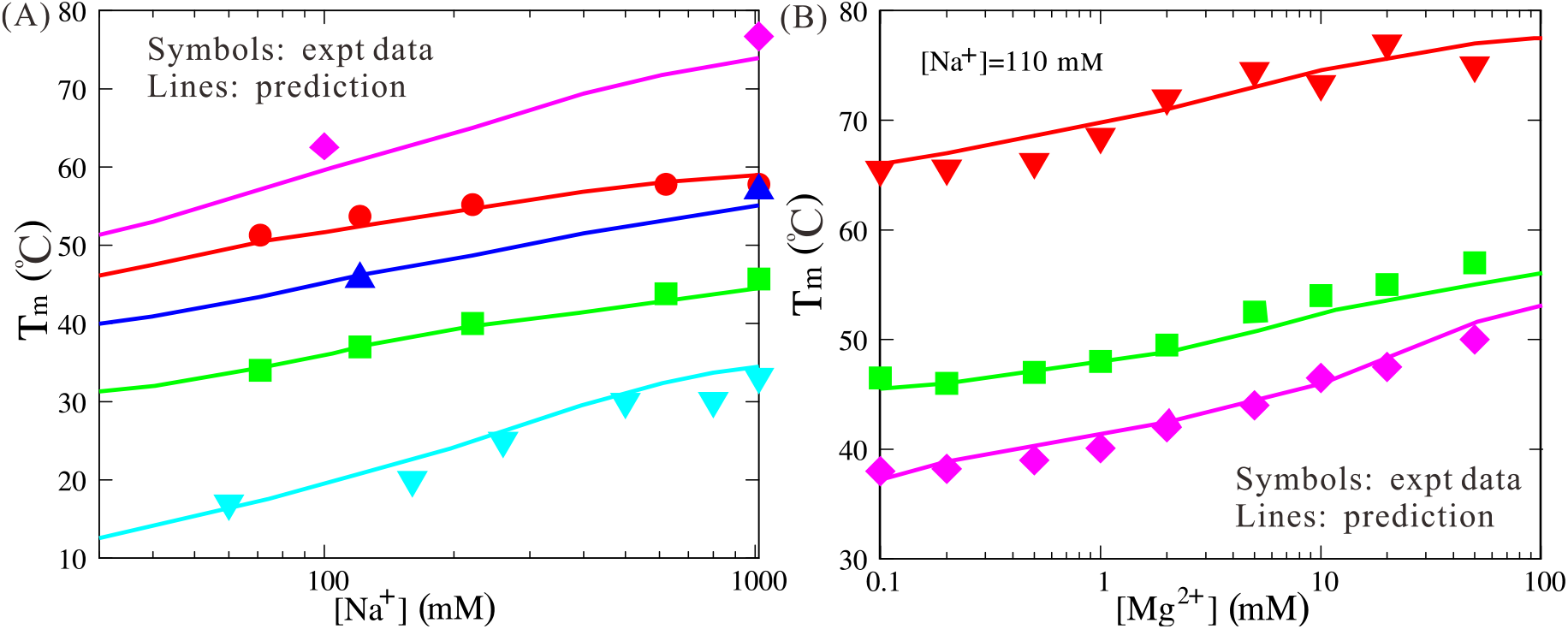
(A) The predicted melting temperatures *T_m_* and corresponding experimental data (65,66,68-70,84) as functions of [Na^+^] for sequences: CCUUGAUAUCAAGG, CGCGCG, CCAUAUGG, AACUAGUU and AAAAAAAUUUUUUU (from top to bottom). (B) The predicted melting temperatures *T_m_* and corresponding experimental data (84) as functions of [Mg^2+^] for sequences: CCUUGAUAUCAAGG, CCAUAUGG and CCAUGG (from top to bottom). Note that the solutions contain 110 mM Na^+^ as background (84).

Second, we examined the stability of dsRNAs in mixed monovalent and divalent ion solutions. As shown in Fig. 7B, for 3 different dsRNAs, the predicted *T_m_*’s are in good accordance with the experimental data over the wide range of [Mg^2+^] (84). Fig. 7B also shows that there are three ranges in *T_m_*-[Mg^2+^] curves: (i) at low [Mg^2+^] (relatively to [Na^+^]), the stability of the dsRNAs is dominated by the background [Na^+^] and *T_m_*’s of the dsRNAs are almost the same with the corresponding pure [Na^+^]; (ii) with the increase of [Mg^2+^], Mg^2+^ ions begin to play a role and **T_m_** increases correspondingly; (iii) when [Mg^2+^] becomes very high (relatively to [Na^+^]), the stability is dominated by Mg^2+^. Furthermore, it is shown that Mg^2+^ is very efficient in stabilizing dsRNAs. Even in the background of 110 mM Na^+^, ~1 mM Mg^2+^ begins to enhance the stability of dsRNAs, and 10 mM Mg^2+^(+110 mM background Na^+^) can achieve the similar stability to 1 M Na^+^ for dsRNAs; see sequences of CCUUGAUAUCAAGG and CCAUAUGG in Figs. 7A and B. This is attributed to the high ionic charge of Mg^2+^ and the consequent efficient role in stabilizing dsRNAs (56-60, 65,81-84).

### Flexibility of dsRNA in ion solutions

DsRNAs generally are rather flexible in ion solutions due to the polymeric nature, and the flexibility is extremely important for their biological functions. Additionally, dsRNA flexibility is highly dependent on solution ion conditions (85-93). In this section, we further employed the present model to examine the flexibility of a 40-bp RNA helix in ion solutions. The sequence of the dsRNA helix is 5’-CGACUCUACGGAAGGGCAUCCUUCGGGCAUCACUACGCGC-3’ with 57% CG content in its central 30-bp segment and the other chain is fully complementary to it, which is selected according to previous study (91). First, we predicted the 3D structures for the dsRNA helix from the sequence at 25°C, and afterwards, we performed further simulations for the dsRNA helix at various ion conditions based on the predicted structures. The enough conformations at equilibrium at each ion condition were used to analyze the salt-dependent flexibility of the dsRNA helix; see Fig. S2 in Supporting Material.

#### Structure fluctuation of a dsRNA in ion solutions

In the following, we first examined the structure fluctuation of the dsRNA helix through calculating end-to-end distance, RMSD variance and root-mean-square fluctuation (RMSF) at different [Na^+^]’s (85). As [Na^+^] increases, the end-to-end distance of the dsRNA helix decreases, e.g., from ~125 Å at 10 mM [Na^+^] to ~90 Å at 1 M [Na^+^], and simultaneously, the variance of end-to-end distance increases; see Figs. 8A and B. This indicates the stronger bending conformations and the higher bending fluctuation for the dsRNA helix at higher ion concentrations, which is attributed to the stronger ion neutralization on P beads charges and consequently the reduced electrostatic repulsion due to bending (86-94). The RMSD variance of the dsRNA helix at different [Na^+^]’s calculated based on the conformation-averaged reference structure also indicates that the dsRNA helix would become more flexible with the increase of [Na^+^]; see Fig. 8C. To examine local structure fluctuation, we further calculated the RMSF of the centers of each base pairs of the dsRNA helix at different [Na^+^]’s. As shown in Fig. 8D, the RMSF increases as [Na^+^] increases from 0.01 M to 1 M, which is because the stronger ion binding and charge neutralization on P beads enable the larger fluctuation of base pairs along the helix (94). Additionally, end effect contributes to an extra increase of RMSF at the two helical ends (94).

**FIGURE 8.**
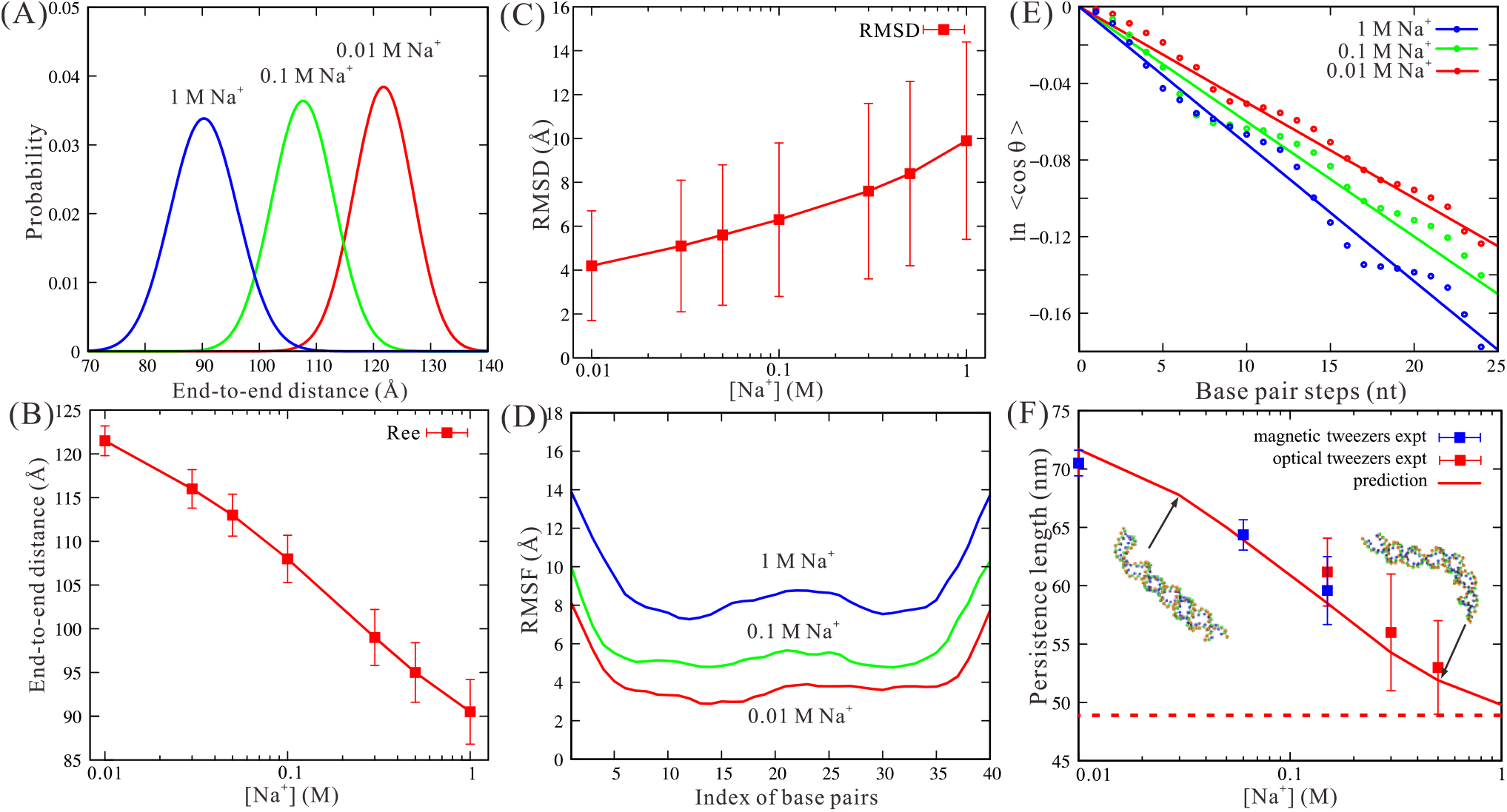
(A) The distributions of end-to-end distance for the 40-bp dsRNA helix at 1 M, 0.1 M and 0.01 M [Na^+^], respectively. (B) The mean end-to-end distances of the 40-bp dsRNA helix as a function of [Na^+^]. Here, the error bars denote the variance for the end-to-end distances. (C) The RMSD of the 40-bp dsRNA helix calculated based on the conformation-averaged reference structure of the respective simulation as a function of [Na^+^]. Here, the error bars denote the variances for the RMSDs. (D) The RMSF of base pairs along the 40-bp dsRNA helix from simulated ensembles at 1 M, 0.1 M and 0.01 M [Na^+^]’s. (E) The fitting curves for *l_p_* through Eq.12 for the 40-bp dsRNA helix at 1 M, 0.1 M and 0.01 M [Na^+^]’s. (F) The predicted persistence length *l_p_* (lines) for the 40-bp dsRNA helix as a function of [Na^+^]. Blue squares: experimental data from magnetic tweezers method (93); red squares: experimental data from optical tweezers method (93). The dashed line in panel (F) shows the predicted *l_p_* with assuming all P beads are electrically neutral.

#### Persistence length of a dsRNA helix in ion solutions

Generally, the flexibility of a polymer can be described by its persistence length *l_p_* (95), and *l_p_* can be calculated by (96):

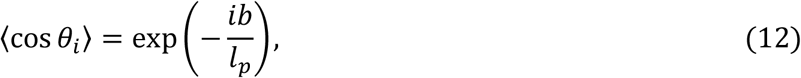

where 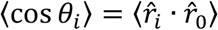 and 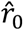 and 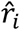 are the first and *i*-th bond direction vectors, respectively. *b* in Eq. 12 is the average bond length. According to Eq. 12, *l_p_* of the dsRNA helix can be obtained through modeling the dsRNA helix as a bead chain composed of the central beads of base pairs; see Fig. 8E. To avoid the end effect (94), the first and last 5 base pairs were excluded in our calculations and the bond vectors are selected as those over every 5 continuous base pairs (91).

As shown in Fig. 8F, the persistence lengths of the 40-bp dsRNA helix at different [Na^+^]’s predicted by the present model are in quantitative agreement with the corresponding experimental data (93). For example, the deviation of *l_p_* between prediction and experiments is less than ~2 nm over the wide range of [Na^+^]. As [Na^+^] increases from 0.01 M to 1 M, *l_p_* of the dsRNA helix decreases from ~70 nm to ~50 nm. This is because more binding ions neutralize the negative P beads charges on the dsRNA helix more strongly and can reduce the electrostatic bending repulsion along the strands more strongly, causing stronger bending flexibility at high [Na^+^].

## Conclusions

Knowledge of the 3D structures and thermodynamic properties of dsRNAs are crucial for understanding their biological functions. In this work, we have further developed our previous CG model by introducing a structure-based electrostatic potential and employed the model to predict 3D structures, stability and flexibility of dsRNAs in monovalent/divalent ion solutions. Our predictions were extensively compared with experimental data, and the following conclusions have been obtained:

1. The present model can well predict 3D structures from sequences for extensive dsRNAs with/without bulge/internal loops in monovalent/divalent ion solutions with overall mean RMSD <3.5 Å, and the involvement of the structure-based electrostatic potential and corresponding experimental ion conditions generally improves the structure predictions with smaller RMSDs for dsRNAs in ion solutions.
2. The present model can make good prediction on the stability for dsRNAs with extensive sequences over wide ranges of monovalent/divalent ion concentrations with mean deviation <2°C, and our analyses show that thermally unfolding pathway of a dsRNA is dependent on its length as well as its sequence.
3. The present model can well capture the salt-dependent flexibility of dsRNAs and the predicted salt-dependent persistence lengths are in good accordance with experiments.

Although our predictions agree well with the extensive experimental data on 3D structure, stability and flexibility of dsRNAs, there are still several limitations in the present model. First, despite that the structure-based electrostatic potential can efficiently capture the effects of monovalent/divalent ions on the structure, stability and flexibility of dsRNAs, the model was not examined for RNAs with more complex structures and the model cannot consider concrete ion distribution and specific ion binding around an RNA. Further development of the present model may need to involve the effect of ions through an implicit-explicit combined treatment for ions (54). Second, the model only involves canonical and wobble base pairing such as A-U, G-C and G-U base pairs, and ignores non-canonical base pairing, which could affect the predictions on the structure and stability of dsRNAs with internal loops. Further involvement of non-canonical base pairing would improve the prediction accuracy of the present model (80). Finally, the present model is a CG model, and it is still necessary to rebuild all-atom structures based on predicted CG ones. Nevertheless, the present model can well predict 3D structures, stability and flexibility of dsRNAs over the wide ranges of monovalent/divalent ion concentrations, and can be a good basis for further development for a predictive model with higher accuracy.

## Acknowledgements

We are grateful to Professors Shi-Jie Chen (University of Missouri), Xiangyun Qiu (The George Washington University), Jian Zhang (Nanjing University) and Wenbing Zhang (Wuhan University) for valuable discussions. This work was supported by grants from the National Science Foundation of China (11575128, 11605125, and 11774272). Parts of the numerical calculation in this work are performed on the super computing system in the Super Computing Center of Wuhan University.

## Author Contributions

Z.J. T., L. J. and Y.Z. S. designed the research; L. J., Y.Z. S and C.J. F. performed the simulation; Z.J. T., L. J., and Y.L. T. analyzed the data; L. J., Y.Z. S. and Z.J. T. wrote the manuscript. All authors discussed the result and reviewed the manuscript.

## Supporting Material

Supporting Material are available at XXX.

## References

1 Watson, J. D. (2008) Molecular biology of the gene. Pearson/Benjamin Cummings.

2 Tinoco, I., and Bustamante, C. (1999) How RNA folds. J. Mol. Biol., 293, 271–281.

3 Li, P. T., Vieregg, J., and Tinoco, I. (2008) How RNA unfolds and refolds. Annu. Rev. Biochem., 77, 77–100.

4 Kuwabara, T., Hsieh, J., Nakashima, K., Taira, K., and Gage, F. H. (2004) A small modulatory dsRNA specifies the fate of adult neural stem cells. Cell, 116, 779–793.

5 Hannon, G. J. (2002) RNA interference. Nature, 418, 244–251.

6 Meister, G., and Tuschl, T. (2004) Mechanisms of gene silencing by double-stranded RNA. Nature, 431, 343–349.

7 Akira, S., and Takeda, K. (2004) Toll-like receptor signalling. Nature reviews immunology, 4, 499–511.

8 Chen, S. J. (2008) RNA folding: conformational statistics, folding kinetics, and ion electrostatics. Annu. Rev. Biophys, 37, 197–214.

9 Mustoe, A. M., Brooks, C. L., and Al-Hashimi, H. M. (2014) Hierarchy of RNA functional dynamics. Annu. Rev. biochem., 83, 441–466.

10 Draper, D. E., Grilley, D., and Soto, A. M. (2005) Ions and RNA folding. Annu. Rev. Biophys. Biomol. Struct., 34, 221–243.

11 Lipfert, J., Doniach, S., Das, R., and Herschlag, D. (2014) Understanding nucleic acid-ion interactions. Annu. Rev. biochem., 83, 813–841.

12 Woodson, S. A. (2005) Metal ions and RNA folding: a highly charged topic with a dynamic future. Curr Opin. Chem. Biol., 9, 104–109.

13 Draper, D. E. (2013) Folding of RNA tertiary structure: linkages between backbone phosphates, ions, and water. Biopolymers, 99, 1105–1113.

14 Koculi, E., Hyeon, C., Thirumalai, D., and Woodson, S. A. (2007) Charge density of divalent metal cations determines RNA stability. J. Am. Chem. Soc., 129, 2676–2682.

15 Rose, P. W., Prlic, A., Altunkaya, A., Bi, C., Bradley, A. R., Christie, C. H., Costanzo, L. D., Duarte, J. M., Dutta, S., Feng, Z., Green, R. K., Goodsell, D. S., Hudson, B., Kalro, T., Lowe, R., Peisach, E., Randle, C., Rose, A. S., Shao, C., Tao, Y. P., Valasatava, Y., Voigt, M., Westbrook, J. D., Woo, J., Yang, H., Young, J. Y., Zardecki, C., Berman, H. M. and Burley, S. K. (2017) The RCSB protein data bank: integrative view of protein, gene and 3D structural information. Nucleic Acids Res., 45, D271–D281.

16 Sim, A. Y., Minary, P., and Levitt, M. (2012) Modeling nucleic acids. Curr. Opin. Struct. Biol., 22, 273–278.

17 Miao, Z. and Westhof, E. (2017) RNA Structure: advances and assessment of 3D structure prediction. Annu. Rev. Biophys., 46, 483–503.

18 Schlick, T. and Pyle, A. M. (2017) Opportunities and challenges in RNA structural modeling and design. Biophys. J. 113, 225–234.

19 Sun, L. Z., Zhang, D., and Chen, S. J., (2017) Theory and modeling of RNA structure and interactions with metal ions and small molecules. Annu. Rev. Biophys., 46, 227–246.

20 Somarowthu, S. (2016) Progress and current challenges in modeling large RNAs. J. Mol. Biol. 428, 736–747.

21 Shi, Y. Z., Wu, Y. Y., Wang, F. H. and Tan, Z. J. (2014) RNA structure prediction: progress and perspective. Chin. Phys. B., 23, 078701.

22 Cragnolini, T., Derreumaux, P. and Pasquali, S. (2015) Ab initio RNA folding. J. Phys. Condens. Matt., 27, 233102.

23 Zhou, H. X. (2014) Theoretical frameworks for multiscale modeling and simulation. Curr. Opin. Struct. Biol., 25, 67–76.

24 Parisien, M. and Major, F. (2008) The MC-Fold and MC-Sym pipeline infers RNA structure from sequence data. Nature., 452, 51–55.

25 Zhao, Y., Huang, Y., Gong, Z., Wang, Y., Man, J. and Xiao, Y. (2012) Automated and fast building of three-dimensional RNA structures. Sci. Rep., 2, 734104.

26 Wang, J., Zhao, Y., Zhu, C., and Xiao, Y. (2015) 3dRNAscore: a distance and torsion angle dependent evaluation function of 3D RNA structures. Nucleic Acids Res., 43, e63.

27 Wang, J., Mao, K., Zhao, Y., Zeng, C., Xiang, J., Zhang, Y. and Xiao, Y. (2017) Optimization of RNA 3D structure prediction using evolutionary restraints of nucleotide-nucleotide interactions from direct coupling analysis. Nucleic Acids Res., 45, 6299–6309.

28 Popenda, M., Szachniuk, M., Antczak, M., Purzycka, K. J., Lukasiak, P., Bartol, N., Blazewicz, J., and Adamiak, R. W. (2012) Automated 3D structure composition for large RNAs. Nucleic Acids Res., 40, e112–e112.

29 Cao, S. and Chen, S. J. (2011) Physics-based de novo prediction of RNA 3D structures. J. Phys. Chem. B., 115, 4216–4226.

30 Xu, X., Zhao, P. and Chen, S. J. (2014) Vfold: a web server for RNA structure and folding thermodynamics prediction. PLoS One., 9, e107504.

31 Das, R. and Baker, D. (2007) Automated de novo prediction of native-like RNA tertiary structures. Proc. Natl. Acad. Sci. USA, 104, 14664–14669.

32 Hyeon, C., and Thirumalai, D. (2011) Capturing the essence of folding and functions of biomolecules using coarse-grained models. Nat. Commun., 2, 487.

33 Jonikas, M. A., Radmer, R. J., Laederach, A., Das, R., Pearlman, S., Herschlag, D. and Altman, R. B. (2009) Coarse-grained modeling of large RNA molecules with knowledge-based potentials and structural filters. RNA, 15, 189–199.

34 Boudard, M., Barth, D., Bernauer, J., Denise, A. and Cohen, J. (2017) GARN2: coarse-grained prediction of 3D structure of large RNA molecules by regret minimization. Bioinformatics, 33, 2479–2486.

35 Kim, N., Laing, C., Elmetwaly, S., Jung, S., Curuksu, J. and Schlick, T. (2014) Graph-based sampling for approximating global helical topologies of RNA. Proc. Natl. Acad. Sci. USA, 111, 4079–4084.

36 Jain, S., and Schlick, T. (2017) F-RAG: Generating Atomic Coordinates from RNA Graphs by Fragment Assembly. J. Mol. Biol., 429, 3587–3605.

37 Zhang, J., Bian, Y., Lin, H., and Wang, W. (2012) RNA fragment modeling with a nucleobase discrete-state model. Phys. Rev. E., 85, 021909.

38 Bian, Y, Zhang, J, Wang, J, Wang, J, Wang, W. (2015) Free energy landscape and multiple folding pathways of an H-Type RNA pseudoknot. PLoS One, 10: e0129089.

39 Li, J., Zhang, J., Wang, J., Li, W. and Wang, W. (2016) Structure prediction of RNA loops with a probabilistic approach. PLoS Comput. Biol., 12, e1005032.

40 Uusitalo, J. J., Ingolfsson, H. I., Marrink, S. J. and Faustino, I. (2017) Martini Coarse-Grained force field: extension to RNA. Biophys. J., 113, 246–256.

41 Sieradzan, A. K., Makowski, M., Augustynowicz, A., and Liwo, A. (2017). A general method for the derivation of the functional forms of the effective energy terms in coarse-grained energy functions of polymers. I. Backbone potentials of coarse-grained polypeptide chains. J. Chem. Phys., 146, 124106.

42 Ding, F., Sharma, S., Chalasani, P., Demidov, V. V., Broude, N. E. and Dokholyan, N. V. (2008) Ab initio RNA folding by discrete molecular dynamics: From structure prediction to folding mechanisms. RNA, 14, 1164–1173.

43 Boniecki, M. J., Lach, G., Dawson, W. K., Tomala, K., Lukasz, P., Soltysinski, T., Rother, K. M. and Bujnicki, J. M. (2016) SimRNA: a coarse-grained method for RNA folding simulations and 3D structure prediction. Nucleic Acids Res., 44, e63.

44 Cragnolini, T., Derreumaux, P. and Pasquali, S. (2013) Coarse-Grained simulations of RNA and DNA duplexes. J. Phys. Chem. B., 117, 8047–8060.

45 Xia, Z., Bell, D. R., Shi, Y. and Ren, P. (2013) RNA 3D structure prediction by using a Coarse-Grained model and experimental data. J. Phys. Chem. B., 117, 3135–3144.

46 Bell, D. R., Cheng, S. Y., Salazar, H., and Ren, P. (2017) Capturing RNA Folding Free Energy with Coarse-Grained Molecular Dynamics Simulations. Scientific Reports, 7, 45812.

47 Denesyuk, N. A. and Thirumalai, D. (2013) Coarse-Grained model for predicting RNA folding thermodynamics. J. Phys. Chem. B., 117, 4901–4911.

48 Hori N., Denesyuk N. A., Thirumalai D. (2016) Salt effects on the thermodynamics of a frameshifting RNA pseudoknot under tension. J. Mol. Biol., 428: 2847–2859.

49 Šulc, P., Romano, F., Ouldridge, T. E., Doye, J. P. K. and Louis, A. A. (2014) A nucleotide-level coarse-grained model of RNA. J. Chem. Phys., 140, 235102.

50 He, Y., Maciejczyk, M., Ołdziej, S., Scheraga, H. A. and Liwo, A. (2013) Mean-field interactions between nucleic-acid-base dipoles can drive the formation of a double helix. Phys. Rev. Lett., 110, 98101.

51 He, Y., Liwo, A. and Scheraga, H. A. (2015) Optimization of a Nucleic Acids united-RESidue 2-Point model (NARES-2P) with a maximum-likelihood approach. J. Chem. Phys., 143, 243111.

52 Manning, G. S. (1978) The molecular theory of polyelectrolyte solutions with applications to the electrostatic properties of polynucleotides. Q. Rev. Biophys., 11, 179–246.

53 Hayes, R. L., Noel, J. K., Mandic, A., Whitford, P. C., Sanbonmatsu, K. Y., Mohanty, U. and Onuchic, J. N. (2015) Generalized manning condensation model captures the RNA ion atmosphere. Phys. Rev. Lett., 114, 258105.

54 Shi, Y. Z., Wang, F. H., Wu, Y. Y. and Tan, Z. J. (2014) A coarse-grained model with implicit salt for RNAs: Predicting 3D structure, stability and salt effect. J. Chem. Phys., 141, 105102.

55 Shi, Y. Z., Jin, L., Wang, F. H., Zhu, X. L. and Tan, Z. J. (2015) Predicting 3D structure, flexibility, and stability of RNA hairpins in monovalent and divalent ion solutions. Biophys. J., 109, 2654–2665.

56 Tan, Z. J. and Chen, S. J. (2005) Electrostatic correlations and fluctuations for ion binding to a finite length polyelectrolyte. J. Chem. Phys., 122, 44903.

57 Tan, Z. J. and Chen, S. J. (2011) Salt contribution to RNA tertiary structure folding stability. Biophys. J., 101, 176–187.

58 Tan, Z. J. and Chen, S. J. (2010) Predicting ion binding properties for RNA tertiary structures. Biophys. J., 99, 1565–1576.

59 Wang, F. H., Wu, Y. Y. and Tan, Z. J. (2013) Salt contribution to the flexibility of single-stranded nucleic acid of finite length. Biopolymers., 99, 370–381.

60 Xi, K., Wang, F., Xiong, G., Zhang, Z. and Tan, Z. J. (2018) Competitive binding of Mg^2^+ and Na+ ions to nucleic acids: from helices to tertiary structures. Biophys. J., 114, 1776–1790.

61 Privalov, P. L. and Crane-Robinson, C. (2018) Translational entropy and DNA duplex stability. Biophys J., 114, 15–20.

62 Cao, S., and Chen, S. J. (2006) Free energy landscapes of RNA/RNA complexes: with applications to snRNA complexes in spliceosomes. J. Mol. Biol., 357, 292–312.

63 Ouldridge, T. E., Louis, A. A. and Doye, J. P. (2010) Extracting bulk properties of self-assembling systems from small simulations. J. Phys. Condens. Matt., 22, 104102.

64 Borer, P. N., Dengler, B., Tinoco, I. J. and Uhlenbeck, O. C. (1974) Stability of ribonucleic acid double-stranded helices. J Mol Biol., 86, 843–853.

65 Tan, Z. J. and Chen, S. J. (2007) RNA Helix Stability in Mixed Na+/Mg^2^+ Solution. Biophys. J. 92, 3615–3632.

66 Xia, T., SantaLucia, J., Burkard, M. E., Kierzek, R., Schroeder, S. J., Jiao, X., Cox, C. and Turner, D. H. (1998) Thermodynamic Parameters for an Expanded Nearest-Neighbor Model for Formation of RNA Duplexes with Watson-Crick Base Pairs. Biochemistry., 37, 14719–14735.

67 Parisien, M., Cruz, J. A., Westhof, E. and Major, F. (2009) New metrics for comparing and assessing discrepancies between RNA 3D structures and models. RNA, 15, 1875–1885.

68 Nakano, S., Fujimoto, M., Hara, H. and Sugimoto, N. (1999) Nucleic acid duplex stability: influence of base composition on cation effects. Nucleic Acids Res., 27, 2957–2965.

69 Hickey, D. R. and Turner, D. H. (1985) Solvent Effects on the Stability of A7U7p. Biochemistry., 24, 2086–2094.

70 Chen, Z. and Znosko, B. M. (2013) Effect of sodium ions on RNA duplex Stability. Biochemistry., 52, 7477–7485.

71 Wang Y, Gong S, Wang Z, Zhang W. (2016) The thermodynamics and kinetics of a nucleotide base pair. J. Chem. Phys., 144: 115101.

72 Shi, Y. Z, Jin, L., Feng, C. J., Tan, Y. L. and Tan, Z. J. (2018) Predicting 3D structure and stability of RNA pseudoknots in monovalent and divalent ion solutions. PLoS Comput. Biol., (in press).

73 Zuker M. (2003) Mfold web server for nucleic acid folding and hybridization prediction. Nucleic Acids Res., 31: 3406–3415.

74 Tomcho, J. C., Tillman, M. R. and Znosko, B. M. (2015) Improved model for predicting the free energy contribution of dinucleotide bulges to RNA duplex stability. Biochemistry, 54, 5290–5296.

75 Crowther, C. V., Jones, L. E., Morelli, J. N., Mastrogiacomo, E. M., Porterfield, C., Kent, J. L. and Serra, M. J. (2016) Influence of two bulge loops on the stability of RNA duplexes. RNA, 23, 217–228.

76 Znosko, B. M., Silvestri, S. B., Volkman, H., Boswell, B. and Serra, M. J. (2002) Thermodynamic parameters for an expanded nearest-neighbor model for the formation of RNA duplexes with single nucleotide bulges. Biochemistry, 41, 10406–10417.

77 Murray, M. H., Hard, J. A. and Znosko, B. M. (2014) Improved model to predict the free energy contribution of trinucleotide bulges to RNA duplex stability. Biochemistry, 53, 3502–3508.

78 Chen, G., Znosko, B. M., Jiao, X. and Turner, D. H. (2004) Factors affecting thermodynamic stabilities of RNA 3 × 3 internal loops. Biochemistry, 43, 12865–12876.

79 Longfellow, C. E., Kierzek, R. and Turner, D. H. (1990) Thermodynamic and spectroscopic study of bulge loops in oligoribonucleotide. Biochemistry, 29, 278.

80 SantaLucia, J., Kierzek, R. and Turner, D. H. (1991) Stabilities of consecutive AC, CC, GG, UC, and UU Mismatches in RNA internal loops: evidence for stable Hydrogen-Bonded UU and CC Pairs. Biochemistry, 30, 8242.

81 Zhang, Z. L., Wu, Y. Y., Xi, K., Sang, J. P., and Tan, Z. J. (2017). Divalent ion-mediated DNA-DNA interactions: A comparative study of triplex and duplex. Biophys. J., 113, 517–528.

82 Denesyuk, N. A., and Thirumalai, D. (2015). How do metal ions direct ribozyme folding? Nature Chem., 7, 793.

83 Leipply, D. and Draper, D. E. (2011) Effects of Mg^2^+ on the free energy landscape for folding a purine riboswitch RNA. Biochemistry, 50, 2790–2799.

84 Serra, M. J., Baird, J. D., Dale, T., Fey, B. L., Retatagos, K. and Westhof, E. (2002) Effects of magnesium ions on the stabilization of RNA oligomers of defined structures. RNA, 8, 307–323.

85 Hagerman, P. J. (1997). Flexibility of RNA. Annu. Rev. Biophys. Biomol. Struct., 26, 139–156.

86 Bao, L., Zhang, X., Jin, L., and Tan Z. J., (2016). Flexibility of nucleic acids: From DNA to RNA. Chin. Phys. B, 25.018703.

87 Li, J., Wijeratne, S. S., Qiu, X., and Kiang, C. H. (2015). DNA under force: mechanics, electrostatics, and hydration. Nanomaterials, 5, 246–267.

88 Sutton, J. L., and Pollack, L., (2015) Tuning RNA flexibility with helix length and junction sequence. Biophys. J., 109(12), 2644–2653.

89 Chen, H., Meisburger, S. P., Pabit, S. A., Sutton, J. L., Webb, W. W., and Pollack, L. (2012). Ionic strength-dependent persistence lengths of single-stranded RNA and DNA. Proc. Natl. Acad. Sci. USA, 109, 799–804.

90 Drozdetski, A. V., Tolokh, I. S., Pollack, L., Baker, N. A., and Onufriev, A. V., (2016) Opposing effects of multivalent ions on the flexibility of DNA and RNA. Phys. Rev. Lett., 117, 028101.

91 Bao, L., Zhang, X., Shi, Y. Z., Wu, Y. Y. and Tan, Z. J. (2017) Understanding the relative flexibility of RNA and DNA duplexes: stretching and twist-stretch coupling. Biophys J., 112, 1094–1104.

92 Zhang, X., Bao, L., Wu, Y. Y., Zhu, X. L. and Tan, Z. J. (2017) Radial distribution function of semiflexible oligomers with stretching flexibility. J. Chem. Phys., 147, 054901.

93 Herrero-Galán, E., Fuentes-Perez, M. E., Carrasco, C., Valpuesta, J. M., Carrascosa, J. L., Moreno-Herrero, F. and Arias-Gonzalez, J. R. (2013) Mechanical identities of RNA and DNA double helices unveiled at the single-molecule level. J Am. Chem. Soc., 135, 122–131.

94 Wu, Y. Y., Bao, L., Zhang, X. and Tan, Z. J. (2015) Flexibility of short DNA helices with finite-length effect: From base pairs to tens of base pairs. J. Chem. Phys., 142, 125103.

95 Kebbekus, P., Draper, D. E. and Hagerman, P. J. (1995) Persistence Length of RNA., Biochemistry., 34, 4354–4357.

96 Ullner, M. and Woodward, C. E. (2002) Orientational correlation function and persistence lengths of flexible polyelectrolytes. Macromolecules., 35, 1437–1445.

